# Evaluating the Sex Dependent Influence of Sarcospan on Cardiometabolic Disease Traits in Mice

**DOI:** 10.1101/2024.03.31.586423

**Authors:** Aida Rahimi Kahmini, Isela C. Valera, Luaye Samarah, Rhiannon Q. Crawford, Salma Elsheikh, Rosemeire M. Kanashiro-Takeuchi, Bolade S. Olateju, Aaron R. Matthews, Nazanin Mohammadipoor, Michelle S. Parvatiyar

## Abstract

Numerous genes including sarcospan (SSPN) have been designated as obesity-susceptibility genes by human genome-wide association studies. Variants in the sarcospan (SSPN) gene locus have been associated with obesity traits with a stronger effect in women. To date this association has not been tested in vivo, therefore, we assessed the susceptibility of young (2 month) global SSPN-deficient (SSPN^-/-^) mice to diet-induced obesity by feeding them high fat diet (HFD) or control diet (CD) for 16 weeks. Anthropometric measurements were used to assess outcomes to the diets including weight change, glucose handling, fat distribution, adipocyte size and effects on cardiac function. To assess the age-dependent impact of SSPN deletion we also compared the response of (13 month) male and female mice to HFD, which were aged by study completion. SSPN deficiency offered modest protection from weight gain in all groups studied, which was not attributable to reduced food consumption. Aging revealed glucose intolerance for SSPN^-/-^ CD mice, which was significant in females. Young female mice had low % Fat and less visceral adipose tissue accumulation that remained relatively unchanged in HFD groups. However, this protection was lost with aging. SSPN^-/-^ mice did not exhibit decrements in cardiac function in response to HFD. However, aged male SSPN^-/-^ CD mice had significantly increased left ventricular mass (LVM) and signs of ventricular remodeling in response to HFD. These studies suggest that SSPN influences phenotype in a sex dependent manner and participates in a network of metabolic genes.

**New & Noteworthy:** In this study the association of the sarcospan protein with human obesity is assessed using in vivo models. Sarcospan-deficient mice of both sexes show an age-dependent influence on adipose tissue biology and glucose handling in response to control and high fat diet. The effect of sarcospan deletion was more pronounced effects in females. Aging reveals susceptibility of SSPN-deficient male mice to increased left ventricular mass.

## INTRODUCTION

Obesity is on the rise throughout the world and contributes to the development of metabolic and cardiovascular diseases. A recent global report estimates that roughly 500 million adults are obese with a body mass index (BMI) of 30 or higher ^1^. Weight gain and obesity is accompanied by accumulation of abdominal fat, which heightens risk of cardiovascular disease and premature death. Adipose tissue releases a multitude of bioactive factors that influences adipocyte homeostasis and insulin resistance. Development of insulin resistance is associated with classical cardiovascular risk factors that include abnormalities in lipid storage, glucose intolerance and hypertension. Cardiovascular tissue is impacted by insulin resistance and decreased glucose uptake, which can undermine normal cardiac function ^2^. In addition, obesity can lead to development of cardiometabolic syndrome (CMS)^3^. CMS is classified as cluster of metabolic dysfunctions including insulin resistance, impaired glucose handling, dyslipidemia, hypertension, and central adiposity ^4^.

It has been established that obesity is highly heritable and influenced by a number of factors including gene interactions, environment and behavior ^5^. The list of genes associated with obesity susceptibility is growing and many of these gene associations remain to be verified. Recent human genome wide association studies (GWAS) studies have labeled the *SSPN* gene, which encodes an integral protein within the dystrophin-glycoprotein complex (DGC) as an obesity susceptibility gene ^6–12^, however in vivo experimental verification is lacking. Pathogenic variants in DGC proteins have been demonstrated to cause various skeletal muscle, heart, and neurological conditions, as reviewed by ^13^ ^14^. Sarcospan (SSPN) is a small tetraspanin-like protein abundantly expressed in striated ^15^ ^16^ and smooth muscle ^17^ but also other tissues including adipose ^18^. The physiological role of the SSPN protein in striated muscle has been previously studied in SSPN-null (SSPN^-/-^) mice ^16, 19, 20^. Several preclinical studies have shown its promise as a therapeutic agent that stabilizes muscle membranes in Duchenne muscular dystrophy (DMD) mice ^19, 21–23^.

Earlier familial GWAS studies reported single nucleotide polymorphisms (SNPs) in the chromosome region 12p11 that segregates with specific traits including increased waist-circumference (WC) and left ventricular mass (LVM) ^24^. The human SSPN gene locus is located at 12p12.1 (NCBI Gene). Variants within the SSPN locus have been primarily associated with increased mid-section adiposity or waist-to-hip ratio (WHR), waist circumference (WC) and WC/WHR) ^25^ cardiometabolic disease traits including hypertension ^26^, metabolic ^10^ ^27^ and cardiovascular risk factors ^10, 24, 28–30^. The genetic association between SSPN expression and increased left ventricular mass may be a secondary effect of altered metabolic state or effect on systolic, diastolic and pulse blood pressures ^26, 31^. These studies suggest that SSPN has an important yet unknown contribution to key predisposing factors that contribute to development of cardiometabolic syndrome (CMS).

While GWAS provides a tool to understand the complex genetics underlying increased susceptibility to obesity, the biological mechanisms relative to specific genetic risk remain poorly defined ^32^. Several studies describe obesity-associated SNPs in the SSPN promoter that alter DNA methylation around the SSPN and subsequently SSPN expression. These studies also reported a strong sex bias for the *SSPN* gene and obesity in women ^18, 33,34^. The current study examines sex differences in white adipose tissue distribution, glucose tolerance and cardiac function in mice lacking the *SSPN* gene. Cardiovascular disease presents differently between sexes and is therefore an important consideration^35^. Since aging influences these parameters, the role of SSPN throughout lifespan was examined. The goal of this study is to provide experimental verification of multiple reports associating SSPN with obesity and increased predisposition to cardiometabolic disease. In addition, this study will assess the potential of the SSPN protein as a novel therapeutic target for treatment of obesity and associated complications. This study, therefore, was designed to provide a comprehensive view of the role of SSPN in humans and provides anthropometric data obtained from young and aged mice of both sexes.

## MATERIALS AND METHODS

### Mouse models

Mice utilized in this study were WT (C57BL6/J) (JAX#000664) and Sspn knock-out mice (JAX#006837) were obtained from Jackson laboratories. The Sspn knock- out mice were developed by the Campbell laboratory ^16^ and donated to Jackson laboratories. In our laboratory the SSPN-deficient (SSPN^-/-^) mice were backcrossed twice with C57BL/6J and heterozygous crosses were utilized to obtain WT controls and SSPN^-/-^ mice. Group housing of mice of both sexes allows mice to participate in normal behavior patterns, although at a cost to measuring individual food consumption. Group housed female mice stimulates and coordinates the estrus cycle in young mice.

### Study Design

In this study 16 groups of mice were assessed. In the young mouse groups the mice were (2 to 2.5 months old at HFD initiation (4 months), therefore, the following groups were approximately 6 – 6.5 months old at final analysis and their total number is listed 1) WT male CD (n= 21), 2) WT female CD (n=12), 3) WT male HFD (n=14), 4) WT female HFD (n=19), 5) SSPN^-/-^ male CD (n= 23), 6) SSPN^-/-^ female CD (n=19), 7) SSPN^-/-^ male HFD (n=11), 8) SSPN^-/-^ female HFD (n=13). In the aged mouse groups were (11-12 months) old at HFD initiation (5 months), therefore, the following groups were approximately 17-18 months old at final analysis and their total number is listed 9) WT male CD (n= 12), 10) WT female CD (n=12), 11) WT male HFD (n=6), 12) WT female HFD (n=6), 13) SSPN^-/-^ male CD (n=15), 14) SSPN^-/-^ female CD (n=15), 15) SSPN^-/-^ male HFD (n=8), 16) SSPN^-/-^ female HFD (n=10).

### High-Fat diet administration

Both male and female WT and SSPN-null mice were fed Envigo high-fat diet (TD.06414) (60% fat content, 5.1 Kcal/g). The approximate fatty acid profile (% 60 of total fat), comprised of 36% saturated, 41% monosaturated, 23% polyunsaturated). Initial data (not shown) was obtained using the Envigo control diet (TD.08806) (10% fat content, 9% sucrose, 3.6 Kcal/g) as recommended by the manufacturer, however it was found incompatible for the SSPN^-/-^ mice that exhibited liver and kidney enlargement after four months on the diet. Instead, the standard chow diet (LabDiet #5001- RHI-E 14) (4.5% crude fat) ad libitum. The duration of the diet regimen for the young mice was four months starting at 2 months of age and for the old mice the diet was extended to five months starting at 12-13 months of age.

### Anthropometric analysis

The mice were fed normal chow diet (CD) and HFD were weighed once a week to assess changes in weight. In addition, food intake was monitored and weighed. After four months of diet(s) the young mice were weighed, and the aged mice were weighed after five months of diet(s). After sacrifice the tissues were collected and heart, liver, and epididymal fat pad weights, and tibia length were reported.

### Tissue histology

After heart, liver, and white adipose tissues were harvested from mice embedded in OCT and immediately flash frozen in liquid N_2_ cooled 2-methylpentane and sections and cut to 7 μm thickness. <u>Hematoxylin & Eosin</u> (H&E) staining was used to visualize of changes in tissue architecture as well as indicate regions of fibrosis as previously described ^19^. <u>Masson’s Trichrome</u> was used to detect collagen infiltration in the tissue sections as previously reported ^23^. <u>Oil Red O staining</u> was used to visualize lipid oil droplets in tissues (Electron Microscopy Sciences Cat #26609-01). <u>Wet mount images of white</u> <u>adipose tissue</u> were obtained after fixation in a 4% PFA solution prior to imaging and cell area and number measurements. Images were captured under identical conditions using a Leica DMI4000B inverted fluorescent microscope equipped using 10X and 20X objectives, and image acquisition software Axiovision Rel 4.5 software (Carl Zeiss, Inc.).

### Glucose tolerance testing

Pre- and post-diet glucose tolerance tests were conducted for all mice in the study. After mice were fasted overnight (∼15 hrs) the response to a glucose bolus 1g/kg administered IP was measured using glucose strips and glucometer (Advanced Glucose Meter (Cat# 08396-5001-75) at 0, 30, 60, 90, 120 min time points post glucose injection. These experiments were performed first thing in the morning The blood sampling was obtained using the tail nick method. Blood glucose readings assessed at 15-, 60-, 120-, and 180-minute time points for males and 15-, 60-, 120-minutes for females.

### EchoMRI imaging

Body composition measurements were taken for mice at the beginning and completion of the study using the EchoMRI system. Measurements were taken of conscious mice and sunflower oil was used as a 100% fat standard to calibrate the system prior to data acquisition. Mice were weighed prior to measurements and data generated were % Fat, % Lean and % Water based upon weight of the mouse.

### DNA Methylation Assay

VAT was collected from mice and flash frozen in liquid N_2_ until further use. The VAT was weighed prior to use and 50 ug of tissue and DNA was extracted using the Quick-DNA^TM^ Miniprep Plus Kit (Zymo) following manufacturer’s instructions, and 50 mg DNA was used to determine DNA-methylation using the 5-mC DNA ELISA Kit (Cat # D5325-A) following the instructions. A methylation standard was included and anything below this value was considered non-methylated.

### Echocardiography

Measurements were recorded of ventricular (LV) size, wall thickness, mass, ventricular and valve function, and Doppler blood flows were obtained using previously described methods ^36–38^. Briefly, mice were anesthetized with 1-2% isoflurane and body temperatures were maintained at 37° C. Hearts were imaged using the parasternal long-axis and short-axis views to determine cardiac morphology and function. Evaluation of diastolic function was performed using the ratio of the LV trans mitral early peak flow (E wave) to late peak flow (A wave) velocity (E/A ratio). All echocardiographic assessments were analyzed offline using the Vevo LAB 2.1.0 software, and Visual Sonics images taken over 3 to 5 consecutive heartbeats.

### Statistics

All values in the text and figures are presented as mean +/- SEM, unless otherwise indicated. Statistical significance was determined using a two-tailed Student’s t-test to compare two relevant groups. Ordinary one-way ANOVA was used when comparisons were made across all groups, followed by Tukey’s multiple comparisons test. *P* values of <0.05 were considered significant.

### Study Approval

The animal studies included in this study were reviewed and approved by the Institutional Animal Care and Use Committee at Florida State University as protocol #IPROTO202200000003.

## RESULTS

### Assessment of Weight Gain by Young and Aged SSPN^-/-^ Mice in Response to High Fat Diet -

To determine whether SSPN has an influence on whole body metabolism - young (2 months) and adult (13 months) old WT and global SSPN-deficient (SSPN^-/-^) male and female mice were fed control (CD) (4.5% fat) or high fat (HFD) (60% fat) diets for a 4- and 5-month periods respectively. At the end of the study young mice were approximately 6 months of age (equivalent to a 20–30-year-old human) whereas adult mice were approximately 18 months of age (equivalent to a 56-year-old human) and considered old by Jackson Laboratories guidelines. For purposes of this study this group will be called aged since they will be “aged” at the conclusion of this study.

Assessments of body weight (BW) measurements are shown for the different groups of mice in Figure 1. Overall, the different groups of mice had similar weights to their controls at baseline. There were however, two exceptions with young SSPN^-/-^ female mice weighing less at baseline than matching WT controls and aged male SSPN^-/-^ mice weighing less than their WT counterparts (Figure 1A and 1E). Since these two groups of SSPN^-/-^ mice weighed significantly less at baseline the % BW change during the diet period provides a better means of comparison (Figure 1 C, D, G, and H). However, the time course of weight changes for male mice is presented in Figure 1A and female mice in Figure 1E. The overall trend in the young mouse groups was that both male and female SSPN^-/-^ mice gained significantly less weight than WT mice during the HFD course (Figure 1A and 1E).

**Figure 1.**
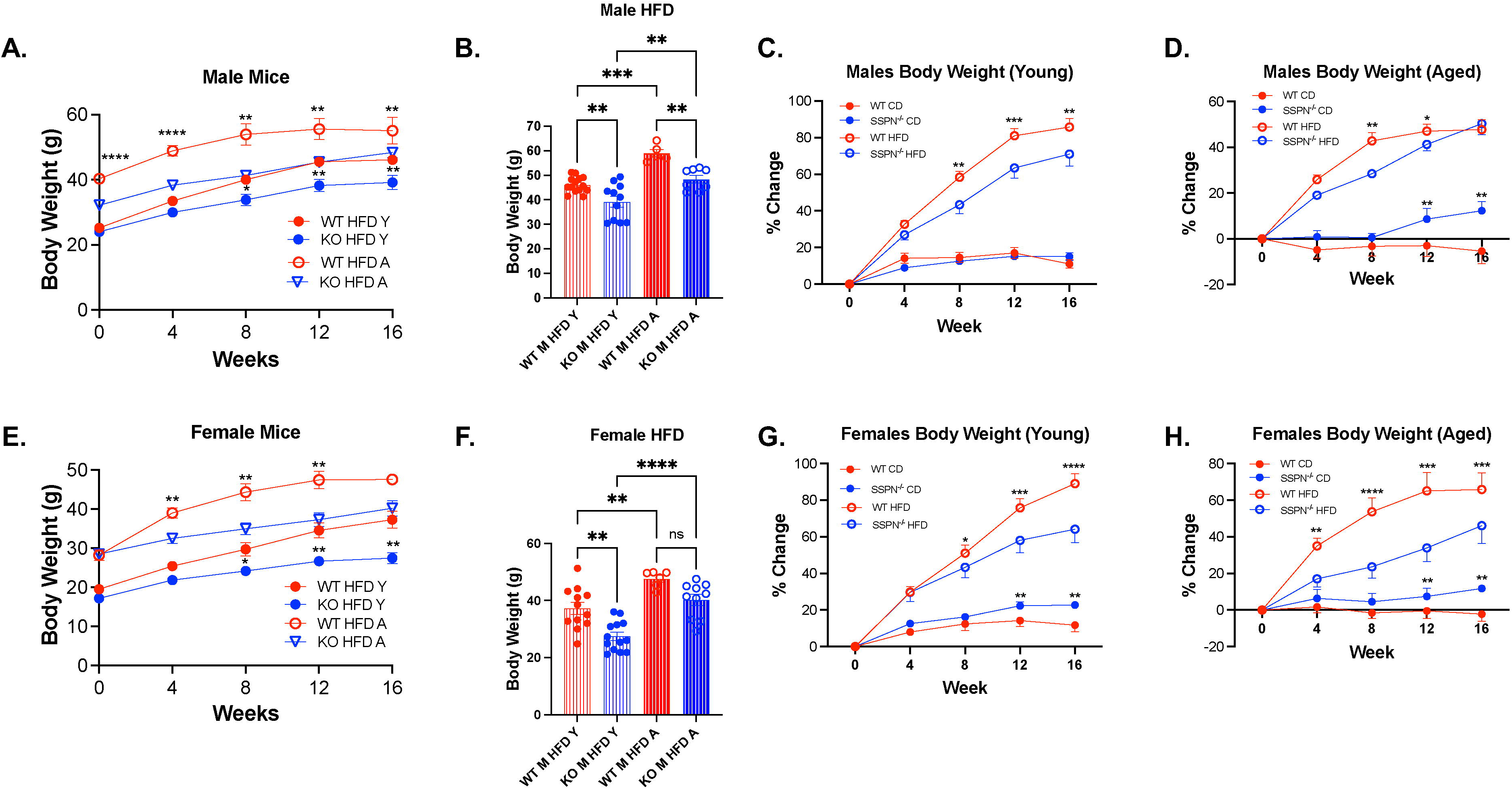
SSPN deficiency is protective against diet-induced obesity. In **(A)** the change in body weight (grams) over 16 weeks high fat diet (HFD) is shown for WT and SSPN- deficient (SSPN^-/-^) male mice with young mice represented by filled circles and aged mice represented by unfilled circles. **(B)** The terminal body weights of the male mice after 16 weeks HFD are shown. **(C)** % change in BW of young male mice **(D)** % change in BW of aged male mice. In **(E)** the change in body weight (grams) over 16 weeks high fat diet (HFD) is shown for WT and SSPN-deficient (SSPN^-/-^) male mice with young mice represented by filled circles and aged mice represented by unfilled circles. **(F)** The terminal body weights of the female mice after 16 weeks HFD are shown. **(G)** % change in BW of young female mice **(H)** % change in BW of aged female mice. Data in panels A, C, D, E, G, H are shown as averages and errors reported as S.E.M. Data in panels B and F are shown as individual values, however bar height represents average value. Data was analyzed by one-way ANOVA followed by Tukey’s post hoc analysis, young males (CD WT (n=6) and SSPN^-/-^ (n=6) and (HFD WT (n=14) and SSPN^-/-^ (n=11)); young females (CD WT (n=6) and SSPN^-/-^ (n=5) and (HFD WT (n=12) and SSPN^-/-^ (n=13)), aged males (CD WT (n=6) and SSPN^-/-^(n=7) and (HFD WT (n=5) and SSPN^-/-^ (n=8), and aged females (CD WT (n=5) and SSPN^-/-^ (n=5) and (HFD WT (n=6) and SSPN^-/-^ (n=10). Data is considered significant if *p*≤0.05 and significance indicated with asterisks, * (<0.05), ** (<0.01), *** (<0.001), **** (<0.0001).

The final weights of the mice in this study are directly compared in Figure 1B after 16 weeks HFD diet mice for males and females in Figure 1F. Upon completion of the CD and HFD the % BW change was compared and both male and female SSPN^-/-^ mice gained significantly less weight than their WT controls (Figure 1A and 1B). After completion of the HFD young WT male mice had a significantly higher 51.58% change in average weight from starting values, whereas HFD young SSPN^-/-^ male mice had an 40.97% increase in average weight (Figure 1C). With aging, the male mice had lower % change than young male mice, however the protection from weight gain during HFD seen in male SSPN^-/-^ mice was largely diminished with % change for WT and SSPN^-/-^ mice of 32.73% and 27.84% respectively (Figure 1D). The WT CD aged male mice appeared to exhibit an age-related decrease in BW with negative % BW change while the SSPN^-/-^ CD mice had a significant increase in BW % change (Figure 1D).

After completion of the HFD studies the young SSPN^-/-^ female mice had significantly lower BW than WT females, however they had lower initial baseline weights (Figure 1F). To assess the difference between the groups, the % change was compared, young WT female HFD mice had a 49.13% increase compared to 39.03% increase for SSPN^-/-^ female mice (Figure 1G). With aging however, the SSPN^-/-^ female HFD mice had a significantly lower % change at 24.14% compared to 43.95% change for WT females (Figure 1H). It was noted in the male young, female young, and female aged CD groups the SSPN^-/-^ mice had significantly higher % change compared to WT mice (Figure 1D, G, and H). All the aged groups of mice had lower % change than the young mouse groups. Multiple factors may be at play including hormonal changes and changes in cellular function including senescence. Food consumption was not significantly different between the HFD young mouse groups (Supplemental Figure 1A). There was however a distinct change with age with all HFD groups consuming fewer grams of chow per day (Supplemental Figure 1B). Interestingly with age the SSPN^-/-^ mice of both sexes appeared to consume approximately one gram/day more than the WT mice though their overall consumption was lower than the young mice.

### Determining the Impact of SSPN Deletion on Glucose Tolerance in Mice –

To assess the effects of SSPN deletion on glucose tolerance, mice were fasted overnight and subjected to glucose tolerance testing. In Figure 2A and Table I the glucose response curves are shown for male mice of all groups in this study. Fasting baseline blood glucose levels were increased in several groups of mice aged SSPN^-/-^ female mice fed CD and HFD and SSPN^-/-^ male mice fed HFD and above 200 mg/dL considered in the diabetic range. One limitation of our study may be use of a human glucometer, which may lead to overestimation of blood glucose in our study. The aged SSPN^-/-^ male mice had the overall highest acute spike in blood glucose at 15 minutes with average values of 559 mg/dL compared to aged WT male mice 463.6 mg/dL. In the graph in Figure 2B the areas under the curve are shown for the male mouse groups. The young male mice had similar AUC values with only a slight increase after 4 months HFD. With age the control male SSPN^-/-^ mice had increased but not significant AUC values compared to WT. Only the WT aged HFD mice exhibited a significant increase in AUC compared to control diet WT mice (Figure 2A).

**Figure 2.**
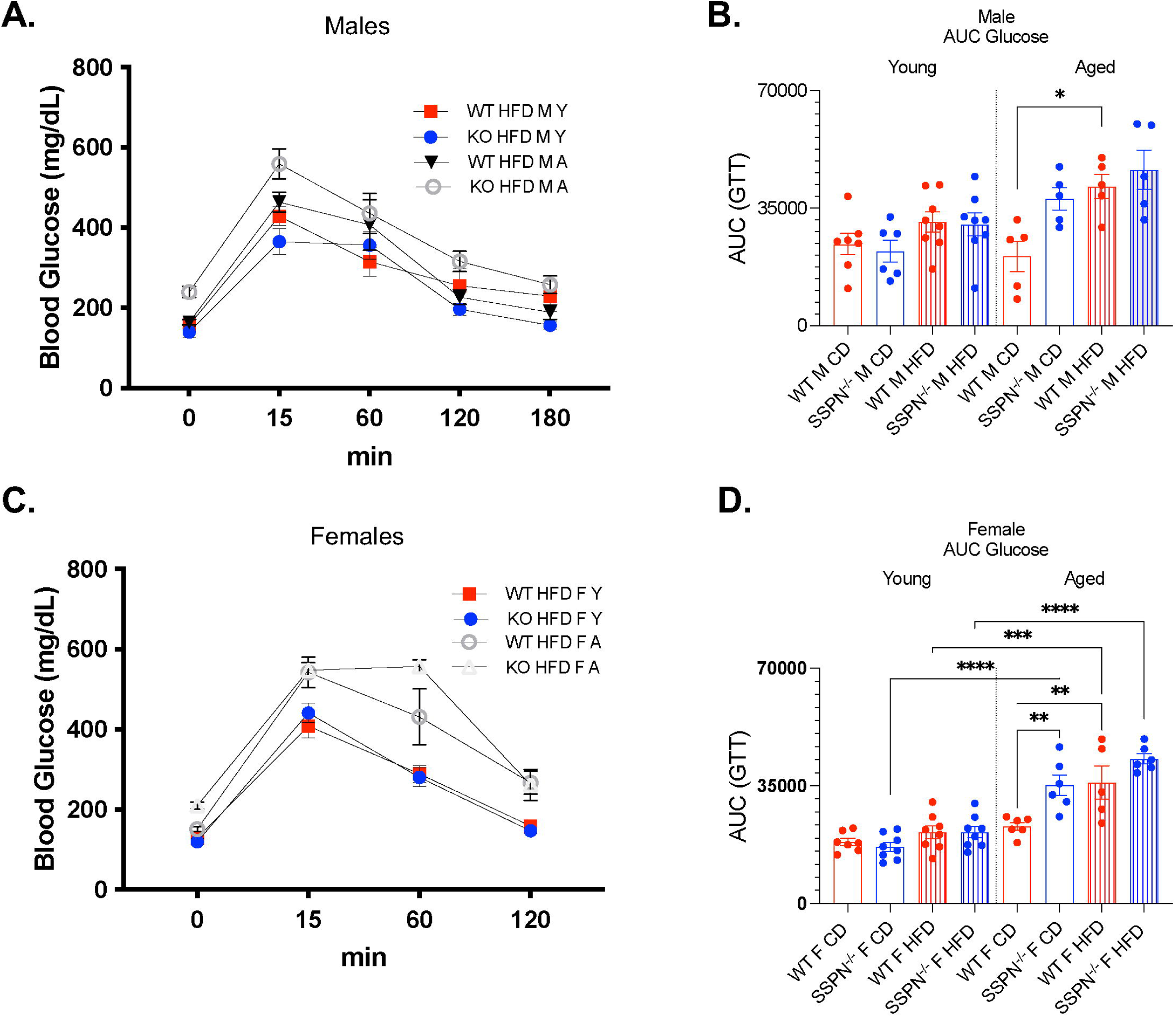
Aging affects glucose tolerance in SSPN-deficient mice. To assess glucose handling in young and aged mice the mice were subjected to glucose tolerance testing. In **(A)** blood glucose values are plotted as a function of time (minutes) for young and aged male WT and SSPN-deficient (SSPN^-/-^) mice after glucose bolus administration from time 0 through 180 minutes. In **(B)** the individual area under the curve (AUC) values for young and aged male mice after glucose bolus administration. In **(C)** alterations in blood glucose are plotted as a function of time (minutes) for young and aged female WT and SSPN-deficient (SSPN^-/-^) mice after glucose bolus administration from time 0 through 120 minutes. **(D)** The graph on the right shows individual values for area under the curve (AUC) for female mice after glucose bolus administration from time 0 through 180 minutes. Data and plotted as averages in A and C and individual values in B and D and error shown as S.E.M. Data was analyzed by one-way ANOVA followed by Tukey’s post hoc analysis and numbers of mice in groups are included in Tables I and II. Significance is indicated with asterisks, * (<0.05), ** (<0.01), *** (<0.001), **** (<0.0001).

**Table I.** Summary of Glucose Tolerance Testing in Control and HFD Young and Aged Male Mice.

| Average Blood Glucose Levels (mg/dL) |  |  |  |  |  |  |  |  |
| --- | --- | --- | --- | --- | --- | --- | --- | --- |
| Time | Young |  |  |  | Aged |  |  |  |
| (minutes) | WT CD<br>(n=7) | SSPN<sup>-/-</sup> CD<br>(n=8) | WT HFD<br>(n=9) | SSPN<sup>-/-</sup> HFD<br>(n=6) | WT CD<br>(n=5) | SSPN<sup>-/-</sup> CD<br>(n=5) | WT HFD<br>(n=5) | SSPN<sup>-/-</sup> HFD<br>(n=5) |
| 0 | 187.2 ± 14.8 | 175.5 ± 9.7 | 156.7 ± 6.3 | 139.7 ± 14.2 | 141 ± 5.8 | 146.6 ± 14.3 | 163.6 ± 6.8 | 239.2 ± 14.1<sup>f,h,k</sup> |
| 15 | 350.2 ± 36.9 | 328.4 ± 29.8 | 428.3 ± 23.6 | 365.2 ± 32.3 | 398.6 ± 43.4 | 409.6 ± 57.4 | 463.6 ± 24.9 | 559 ± 37.4<sup>f</sup> |
| 60 | 246.3 ± 20.2 | 213.2 ± 12.2 | 314.9 ± 36.3 | 356.4 ± 34.7 | 222.6 ± 41.3 | 382.8 ± 72.6 | 406.8 ± 62.9 | 435.4 ± 50.4 |
| 120 | 207.2 ± 21.7 | 217.4 ± 19.7 | 255.2 ± 42.6 | 196.3 ± 14.9 | 150.8 ± 25.4 | 274.6 ± 44.3 | 226.4 ± 18.2 | 315.4 ± 25.4 |
| 180 | 166.5 ± 12.4 | 159 ± 10.2 | 228.7 ± 38.5 | 156.2 ± 9.3 | 116 ± 26.1 | 172.8 ± 19.2 | 189 ± 18 | 257.6 ± 22.2 |

Abbreviations: A (aged), CD (control diet), HFD (high fat diet), M (male), SSPN<sup>-/-</sup> (sarcospan-deficient), WT (wild type), Y (young)
One-way ANOVA was used to compare differences between groups. Superscript designations are:
<sup>a</sup>, $p < 0.05$ - WT (young vs aged) CD mice; <sup>b</sup>, $p < 0.05$ - SSPN<sup>-/-</sup> (young vs aged) CD mice; <sup>c</sup>, $p < 0.05$ - WT and SSPN<sup>-/-</sup> (young) CD mice; <sup>d</sup>, $p < 0.05$ - WT and SSPN<sup>-/-</sup> (young) HFD mice; <sup>e</sup>, $p < 0.05$ - WT (young vs aged) HFD mice; <sup>f</sup>, $p < 0.05$ - SSPN<sup>-/-</sup> (young vs aged) HFD mice; <sup>g</sup>, $p < 0.05$ - WT and SSPN<sup>-/-</sup> (aged) CD mice; <sup>h</sup>, $p < 0.05$ - WT and SSPN<sup>-/-</sup> (aged) HFD mice; <sup>i</sup>, $p < 0.05$ - SSPN<sup>-/-</sup> (aged) CD vs HFD mice; <sup>j</sup>, $p < 0.05$ - WT aged CD vs HFD mice; <sup>k</sup>, $p < 0.05$ - SSPN<sup>-/-</sup> CD vs HFD (aged) mice; <sup>l</sup>, $p < 0.05$ - WT CD vs HFD (aged) mice;
Significant Groups: Time 0m: SSPN<sup>-/-</sup> HFD male (young vs aged), $p < 0.00001$ (\*\*\*\*), Time 15m: SSPN<sup>-/-</sup> HFD male (young vs aged), $p < 0.00001$ (\*\*), SSPN<sup>-/-</sup> CD male

Comparison of the female groups is shown in Figure 2C and Table II, and in the glucose response curve graph (Figure 2D) the aged HFD SSPN^-/-^ mice exhibited sustained elevation of blood glucose for up to 60 mins peaking at 556.7 mg/dL. The young female mice showed a similar trend as male mice, however AUC values were slightly lower. With aging however, significant differences emerged between the groups. The AUC values of aged female SSPN^-/-^ control diet mice were significantly greater compared to both groups of young SSPN^-/-^ mice and WT female HFD mice (Figure 2B). In the aged WT HFD females, a significant increase in AUC was observed when compared to aged CD WT females and young HFD females (Figure 2D).

**Table II.** Summary of Glucose Tolerance Testing in Control and HFD Young and Aged Female Mice.

| Average Blood Glucose Levels (mg/dL) |  |  |  |  |  |  |  |  |
| --- | --- | --- | --- | --- | --- | --- | --- | --- |
| Time | Young |  |  |  | Aged |  |  |  |
| (minutes) | WT CD<br>(n=6) | SSPN<sup>-/-</sup> CD<br>(n=6) | WT HFD<br>(n=8) | SSPN<sup>-/-</sup> HFD<br>(n=8) | WT CD<br>(n=6) | SSPN<sup>-/-</sup> CD<br>(n=6) | WT HFD<br>(n=5) | SSPN<sup>-/-</sup> HFD<br>(n=6) |
| 0 | 151.7 ± 6.8 | 131.5 ± 8.9 | 129.7 ± 5.2 | 119.7 ± 7.9 | 143.5 ± 4.3<sup>a</sup> | 226.2 ± 9.9<sup>b</sup> | 151.2 ± 5.6 | 209.9 ± 8.3<sup>f,h</sup> |
| 15 | 363.9 ± 53.2 | 399.7 ± 45.1 | 408.7 ± 29.9 | 441.3 ± 24.1 | 547.2 ± 33.7 | 472.2 ± 41.9<sup>g</sup> | 542.4 ± 38.1 | 547.7 ± 18.6 |
| 60 | 205.2 ± 14.3 | 191.9 ± 17.2 | 288.9 ± 20.9 | 280.5 ± 23.1 | 231.9 ± 11 | 457.7 ± 42.6<sup>b,g</sup> | 430.8 ± 70.1<sup>e,j</sup> | 556.7 ± 17.2<sup>f</sup> |
| 120 | 175 ± 15.2 | 179.7 ± 12.8 | 158.9 ± 10.4 | 146.9 ± 6.5 | 175.7 ± 10 | 245.9 ± 8.9 | 267 ± 28.5<sup>e,j</sup> | 261.5 ± 38.9<sup>f</sup> |

Abbreviations: A (aged), CD (control diet), HFD (high fat diet), M (male), SSPN<sup>-/-</sup> (Sarcospan-deficient), WT (wild type), Y (young)
One-way ANOVA was used to compare differences between groups. Superscript designations are:
<sup>a</sup>, $p < 0.05$ - WT (young vs aged) CD mice; <sup>b</sup>, $p < 0.05$ - SSPN<sup>-/-</sup> (young vs aged) CD mice; <sup>c</sup>, $p < 0.05$ - WT vs SSPN<sup>-/-</sup> (young) CD mice; <sup>d</sup>, $p < 0.05$ - WT vs SSPN<sup>-/-</sup> (young) HFD mice; <sup>e</sup>, $p < 0.05$ - WT (young vs aged) HFD mice; <sup>f</sup>, $p < 0.05$ - SSPN<sup>-/-</sup> (young vs aged) HFD mice; <sup>g</sup>, $p < 0.05$ - WT vs SSPN<sup>-/-</sup> (aged) CD mice; <sup>h</sup>, $p < 0.05$ - WT vs SSPN<sup>-/-</sup> (aged) HFD mice <sup>i</sup>, $p < 0.05$ - SSPN<sup>-/-</sup> CD vs HFD (aged) mice; <sup>j</sup>, $p < 0.05$ - WT CD vs HFD (aged) mice.
Significant Groups: Time 0: CD SSPN<sup>-/-</sup> (young vs aged)(\*\*\*\*), HFD SSPN<sup>-/-</sup> (aged)(\*\*\*\*), CD WT vs SSPN<sup>-/-</sup> (aged), HFD WT vs SSPN<sup>-/-</sup> (aged)(\*\*\*), Time 15: CD WT (young vs aged)(\*), Time 60: CD WT vs KO (aged)(\*\*\*\*), WT CD vs HFD (aged)(\*\*), Time 120: HFD WT (young vs aged)(\*\*), HFD SSPN<sup>-/-</sup> (young vs aged)(\*\*\*), WT CD vs HFD (aged)(\*).

### Body Composition Measurements to Assess the Impact of Sarcospan on Adipose Distribution -

Body composition measurements by EchoMRI were performed to better understand the role of SSPN in adipose tissue distribution and % body mass. The data was compiled for male mice in Table III and female mice in Table IV. Interestingly when comparing CD groups female SSPN^-/-^ mice trended towards lower % fat while CD male SSPN^-/-^ had significantly lower % fat. After HFD only SSPN^-/-^ males had significantly lower % fat than HFD WT males – however baseline differences may account for this (Figure 3A). Aging had a different influence on CD WT of both sexes with a substantial increase in % fat, while CD SSPN^-/-^ males exhibited negligible increases. The % lean values were lowest in mice which had the greatest increase in % fat (Figure 3C and 3D). Tissue weights after HFD for young mice are reported in (Figure 3E) and for aged mice in (Figure 3F) and provide information concerning changes in organ mass of the mice in this study. In males the visceral adipose tissue (VAT) mass was slightly decreased in the young SSPN^-/-^ mice, however trended higher in the aged SSPN^-/-^ mice. In males no significant differences were seen in liver weights after HFD in the young or aged mice, though liver weights in young SSPN^-/-^ mice trended lower.

**Figure 3.**
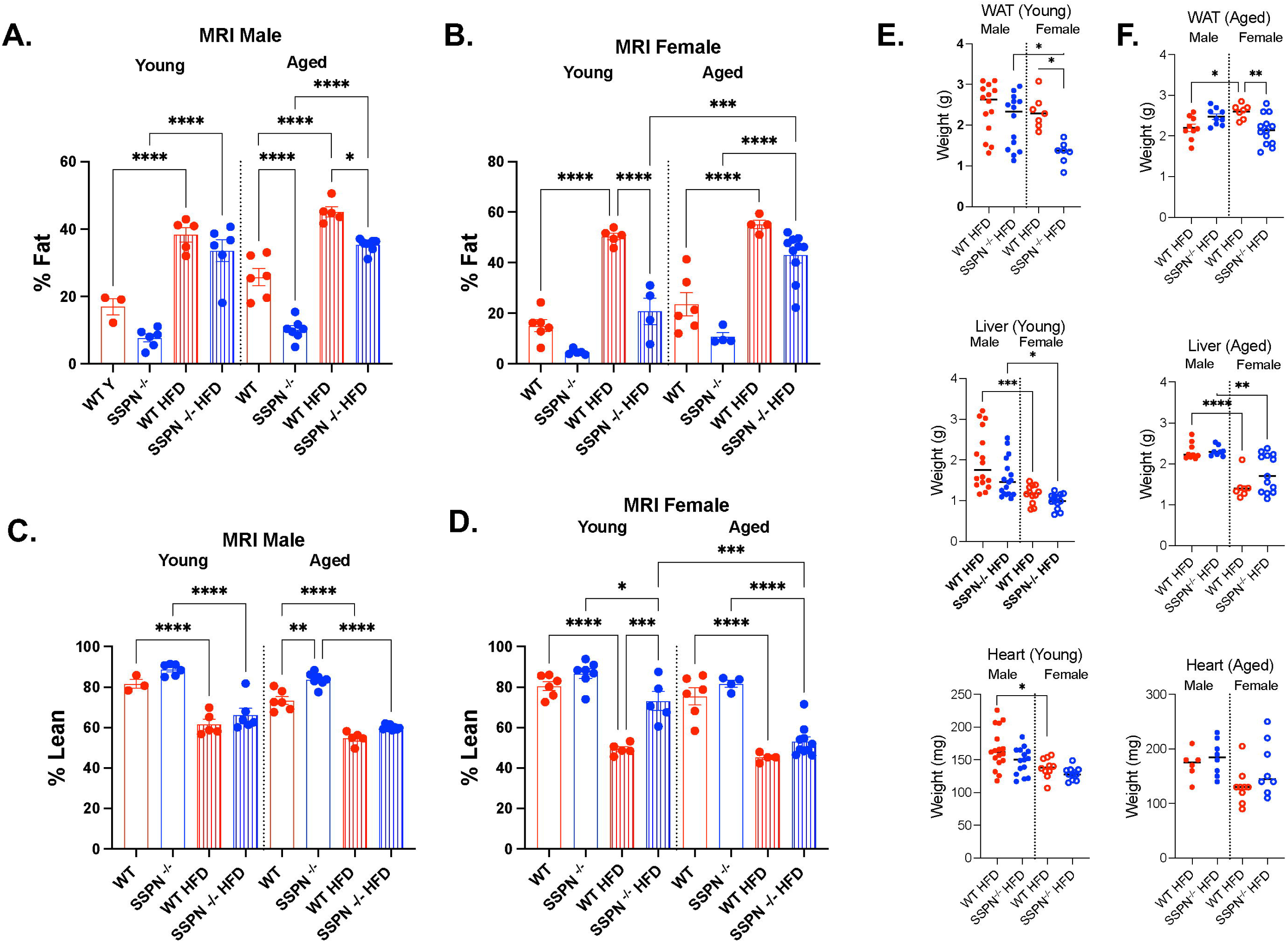
SSPN deficiency causes distinct alterations in white adipose tissue distribution after high-fat diet. EchoMRI was used to assess whole body composition of mice after 12 weeks of either control (CD) or high fat diet (HFD). **(A)** The % Fat measurements from male young and aged WT and SSPN^-/-^ mice are shown and in **(B)** the % Fat measurements from female young and aged WT and SSPN^-/-^ mice are shown. In **(C)** the % Lean measurements from male young and aged WT and SSPN^-/-^ mice are shown and in **(D)** the % Lean measurements from female young and aged WT and SSPN^-/-^ mice are shown. Weights of visceral white adipose tissue (WAT), liver, and heart are shown after CD or HFD in **(E)** for young male and female mice and **(F)** for aged male and female mice. Data was analyzed by one-way ANOVA followed by Tukey’s post hoc analysis. Numbers of mice used in this study are included in Tables III and IV. Data is considered significant if *p*≤0.05 and significance indicated with asterisks, * (<0.05), ** (<0.01), *** (<0.001), **** (<0.0001).

**Table III.** Summary of Body Composition Data in Control and HFD Young and Aged Male Mice.

| EchoMRI Values (Males) |  |  |  |  |  |  |  |  |
| --- | --- | --- | --- | --- | --- | --- | --- | --- |
| Groups | Young |  |  |  | Aged |  |  |  |
|  | WT CD<br>(n=3) | SSPN<sup>-/-</sup> CD<br>(n=6) | WT HFD<br>(n=5) | SSPN<sup>-/-</sup> HFD<br>(n=6) | WT CD<br>(n=6) | SSPN<sup>-/-</sup> CD<br>(n=7) | WT HFD<br>(n=5) | SSPN<sup>-/-</sup> HFD<br>(n=8) |
| Total Fat Percentage (%) | 17 ± 2.4 | 7.7 ± 1.2 | 38.4 ± 2.2<sup>a</sup> | 33.6 ± 3.3<sup>b</sup> | 25.7 ± 2.5 | 10.1 ± 1.2<sup>c</sup> | 45.2 ± 1.5<sup>d</sup> | 35.3 ± 0.7<sup>e,f</sup> |
| Total Lean Percentage (%) | 81.7 ± 2.2 | 88.7 ± 1.2 | 61.7 ± 2.5<sup>a</sup> | 66.3 ± 3.3<sup>b</sup> | 73.4 ± 2.0 | 83.7 ± 1.3<sup>c</sup> | 54.8 ± 1.4<sup>d</sup> | 60.0 ± 0.4<sup>e</sup> |

Abbreviations: CD (control diet), M (male), HFD (high fat diet), SSPN<sup>-/-</sup> (SSPN-deficient), WT (wild type)
One-way ANOVA was used to compare differences between groups. Superscript designations are: <sup>a</sup>, *p*<0.05 – WT CD vs HFD (young) mice; <sup>b</sup>, *p*<0.05 - SSPN<sup>-/-</sup> CD vs HFD (young) mice; <sup>c</sup>, *p*<0.05 - WT vs SSPN<sup>-/-</sup> (aged) CD mice; <sup>d</sup>, *p*<0.05 – WT CD vs HFD (aged) mice, <sup>e</sup>, *p*<0.05 – SSPN<sup>-/-</sup> CD vs HFD (aged) mice; <sup>f</sup>, *p*<0.05 – WT vs SSPN<sup>-/-</sup> HFD (aged) mice.
Significant Groups: Fat (%) WT CD vs HFD (young)(\*\*\*\*), SSPN<sup>-/-</sup> CD vs HFD (young)(\*\*\*\*), CD WT vs SSPN<sup>-/-</sup> (aged)(\*\*\*\*), WT CD vs HFD (aged)(\*\*\*\*), vs SSPN<sup>-/-</sup> CD vs HFD(aged)(\*\*\*\*), WT HFD vs SSPN<sup>-/-</sup> HFD (aged)(\*). Lean (%) WT CD vs HFD (young)(\*\*\*\*), SSPN<sup>-/-</sup> CD vs HFD (young)(\*\*\*\*), CD WT vs SSPN<sup>-/-</sup> (aged)(\*\*), WT CD vs HFD (aged)(\*\*\*\*), vs SSPN<sup>-/-</sup> CD vs HFD(aged)(\*\*\*\*).

**Table IV.** Summary of Body Composition Data in Control and HFD Young and Aged Female Mice.

| EchoMRI Values (Females) |  |  |  |  |  |  |  |  |
| --- | --- | --- | --- | --- | --- | --- | --- | --- |
|  | Young |  |  |  | Aged |  |  |  |
| Groups | WT CD<br>(n=6) | SSPN<sup>-/-</sup> CD<br>(n=7) | WT HFD<br>(n=5) | SSPN<sup>-/-</sup> HFD<br>(n=5) | WT CD<br>(n=6) | SSPN<sup>-/-</sup> CD<br>(n=4) | WT HFD<br>(n=4) | SSPN<sup>-/-</sup> HFD<br>(n=10) |
| Total Fat Percentage (%) | 15.1 ± 2.4 | 4.6 ± 0.5 | 50.4 ± 1.4<sup>a</sup> | 24.4 ± 5.4<sup>b</sup> | 23.5 ± 4.6 | 10.7 ± 1.6<sup>g</sup> | 55.2 ± 1.7<sup>d</sup> | 43.0 ± 3.0<sup>c,e,f</sup> |
| Total Lean Percentage (%) | 80.6 ± 2.2 | 86.7 ± 2.4 | 49.3 ± 1.3<sup>a</sup> | 73.1 ± 4.7<sup>b</sup> | 75.4 ± 4.2 | 81.6 ± 1.7<sup>g</sup> | 45.2 ± 1.1<sup>d</sup> | 53.1 ± 2.4<sup>c,e,f</sup> |

Abbreviations: CD (control diet), M (male), HFD (high fat diet), SSPN<sup>-/-</sup> (SSPN-deficient), WT (wild type)
One-way ANOVA was used to compare differences between groups. Superscript designations are: <sup>a</sup>, *p*<0.05 – WT CD vs HFD (young) mice; <sup>b</sup>, *p*<0.05 - SSPN<sup>-/-</sup> CD vs HFD (young) mice; <sup>c</sup>, *p*<0.05 – SSPN<sup>-/-</sup> HFD (young vs aged) mice; <sup>d</sup>, *p*<0.05 – WT CD vs HFD (aged) mice, <sup>e</sup>, *p*<0.05 – SSPN<sup>-/-</sup> CD vs HFD (aged) mice; <sup>f</sup>, *p*<0.05 – WT HFD vs SSPN<sup>-/-</sup> HFD (aged) mice.
Significant Groups:
Fat (%): WT CD vs HFD (young)(\*\*\*\*), SSPN<sup>-/-</sup> CD vs HFD (young)(\*\*), SSPN<sup>-/-</sup> HFD (young vs aged)(\*\*), WT CD vs HFD (aged)(\*\*\*\*), SSPN<sup>-/-</sup> CD vs HFD (aged)(\*\*\*\*), WT HFD vs SSPN<sup>-/-</sup> HFD (young)(\*\*\*). Lean (%): WT CD vs HFD (young)(\*\*\*\*), SSPN<sup>-/-</sup> CD vs HFD (young)(\*), SSPN<sup>-/-</sup> HFD (young vs aged)(\*\*\*), WT CD vs HFD (aged)(\*\*\*\*), WT HFD vs SSPN<sup>-/-</sup> HFD(young)(\*\*\*).

The most striking findings by EchoMRI were found in the aged SSPN^-/-^ female HFD mice as they had a significant increase in % fat compared to young SSPN^-/-^ female HFD mice (Figure 3B). Furthermore, young SSPN^-/-^ female mice did not have a significant increase in % fat after 16 weeks HFD while WT female mice did (Figure 3B). Aging did not substantially impact other HFD groups as both young and aged mice had similar increases in % fat (Figures 3A and 3B). Regarding % Lean measurements HFD female mice of both genotypes had lower % lean mass than CD controls, inversely proportional to their % Fat values. Overall young SSPN^-/-^ females were largely protected from obesity (low % Fat and preserved % Lean mass), whereas with age this protection was lost and aged SSPN^-/-^ females had significantly higher % fat and a significant reduction in % Lean mass (Figure 3B and 3D). Examination of tissue weights after HFD revealed that young female SSPN^-/-^ mice had significantly lower VAT mass values compared to young WT females (Figure 3E). This trend of lower VAT mass was also seen in aged female SSPN^-/-^, although VAT weights were overall higher than in the young mice (Figure 3F). Height and liver weights, however, were not different between WT and SSPN^-/-^ females in either young or aged groups (Figure 3E and 3F). Therefore, these organs did not contribute to the changes in overall weight of the mice in this study.

### Assessments of Obesogenic Diet and Sarcospan Deletion on Adipocyte Size -

The VAT from each group of mice was further assessed to determine whether SSPN deletion influenced fat storage in VAT under control and obesogenic conditions. In Figure 4A analysis of wet mount images of VAT obtained from young WT HFD male mice revealed adipocytes of varying sizes, with smaller adipocytes indicating either dying or newly formed adipocytes. The VAT obtained from the young SSPN^-/-^ HFD male mice showed larger more uniformly sized adipocytes (Figure 4A). The VAT images from young female WT and SSPN^-/-^ looked largely similar (Figure 4A). Among the young male mouse groups only SSPN^-/-^ HFD mice exhibited a significant increase in adipocyte diameter compared to matching CD mice increasing from 72.27 mm to 96.58 mm (Figure 4C). For females, it differed with young female mice WT HFD having significantly larger adipocyte diameters compared to those found in WT CD VAT (Figure 4D). In this case, WT HFD female mice had significantly larger average adipocyte diameter of 101.5 mm compared to SSPN^-/-^ HFD female mice with 67.10 mm (Figure 4D).

**Figure 4.**
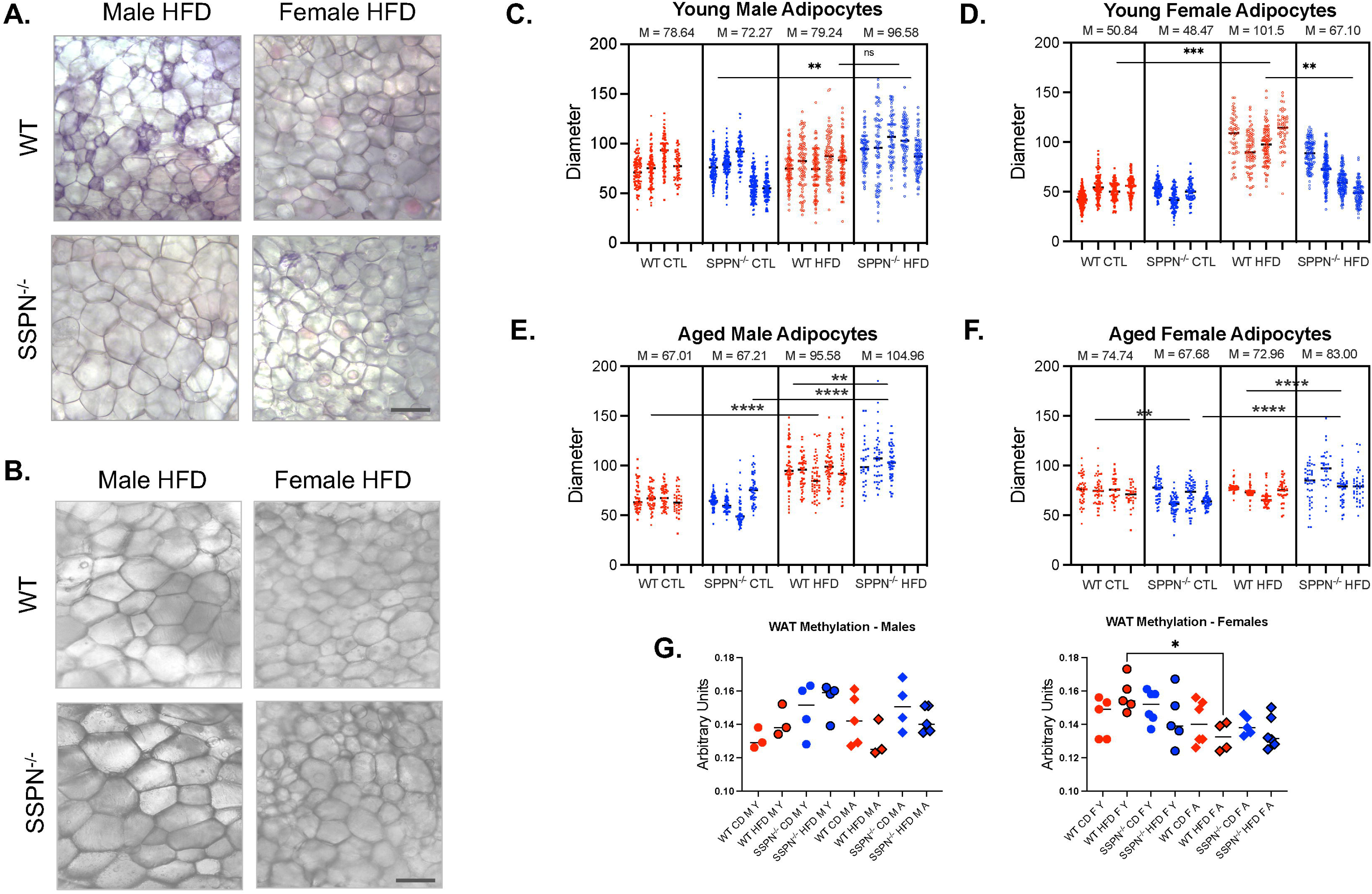
Adipocyte size is altered in some SSPN-deficient mouse groups after high-fat diet. In **(A)** wet mount images of visceral white adipose tissue (VAT) are obtained from young WT and SSPN-deficient (SSPN^-/-^) male and female mice and **(B)** VAT images from aged WT and SSPN-deficient (SSPN^-/-^) male and female mice. Adipocyte area measurements (pixels)^2^ were made using 10X images of wet mount white adipose tissue from WT and SSPN^-/-^ mice. Adipocyte diameters were calculated using d = 2r from area measurements and shown in mm in **(C)** for young males, **(D)** for young females, **(E)** for aged males and **(F)** for aged females. Adipocyte diameters were shown for each individual mouse (n=3-5) and mean values reported above the graphs in C, D, E, F. In **(G)** global DNA methylation levels were assessed using the 5-mC DNA method to determine differences in VAT methylation levels. Images shown are 20X, and Bar = 50 μm. Individual values are shown, and statistics were calculated using one-way ANOVA followed by the Tukey’s multiple comparisons post hoc test and *p* ≥ 0.05 considered significant and errors reported as ± S.E.M.

In the aged male mouse groups, the VAT looked similar between WT and SSPN^-/-^ with larger adipocytes in the HFD groups (Figure 4B). The VAT from the HFD female mice was noticeably smaller (Figure 4B). Measurements of adipocytes showed that male mice of both genotypes had a significant increase in adipocyte hypertrophy, with SSPN^-/-^ adipocyte average diameter of 104.96 mm that was significantly larger than WT adipocytes at 95.58 mm after HFD (Figure 4B). Adipocyte measurements from female VAT indicated smaller adipocytes in SSPN^-/-^ compared to WT CD mice. After HFD however, adipocytes from female SSPN^-/-^ mice were larger than WT with average diameters of 83.00 mm to 72.96 mm respectively (Figure 4F).

Global VAT DNA methylation was assessed to determine whether SSPN deficiency influences this epigenetic modification as reported in studies examining SSPN variants in the promoter region (Figure 4G). The only significant difference between groups was found in female mice with a reduction of VAT DNA methylation in aged WT HFD mice compared to young WT HFD mice. Male showed the same trend in VAT DNA methylation; however, the difference was not significant.

### Evaluating the Impact of SSPN Deletion and Diet-Induced Obesity on Cardiac Function -

Cardiac function was assessed in both young and aged WT and SSPN^-/-^ mice after exposure to metabolic stress. Several anthropometric traits, including increased mid-section adiposity were altered in SSPN^-/-^ mice that serve as predisposing factors for cardiometabolic disease development. In Figure 5A histological evaluation of cardiac tissue from the young HFD mice is shown. The H&E staining revealed minimal fibrotic changes after HFD, however there was a noticeable increase in cellularity (purple stained nuclei) in SSPN^-/-^ hearts of both sexes and male WT hearts. This may be an indication of increased immune cell infiltration when these mice are exposed to obesogenic conditions. In Figure 5B H&E-stained aged male heart tissue showed increased interstitial and perivascular fibrosis. Several SSPN^-/-^ young and aged male hearts had patchy fibrosis, but this was not seen in all the hearts.

**Figure 5.**
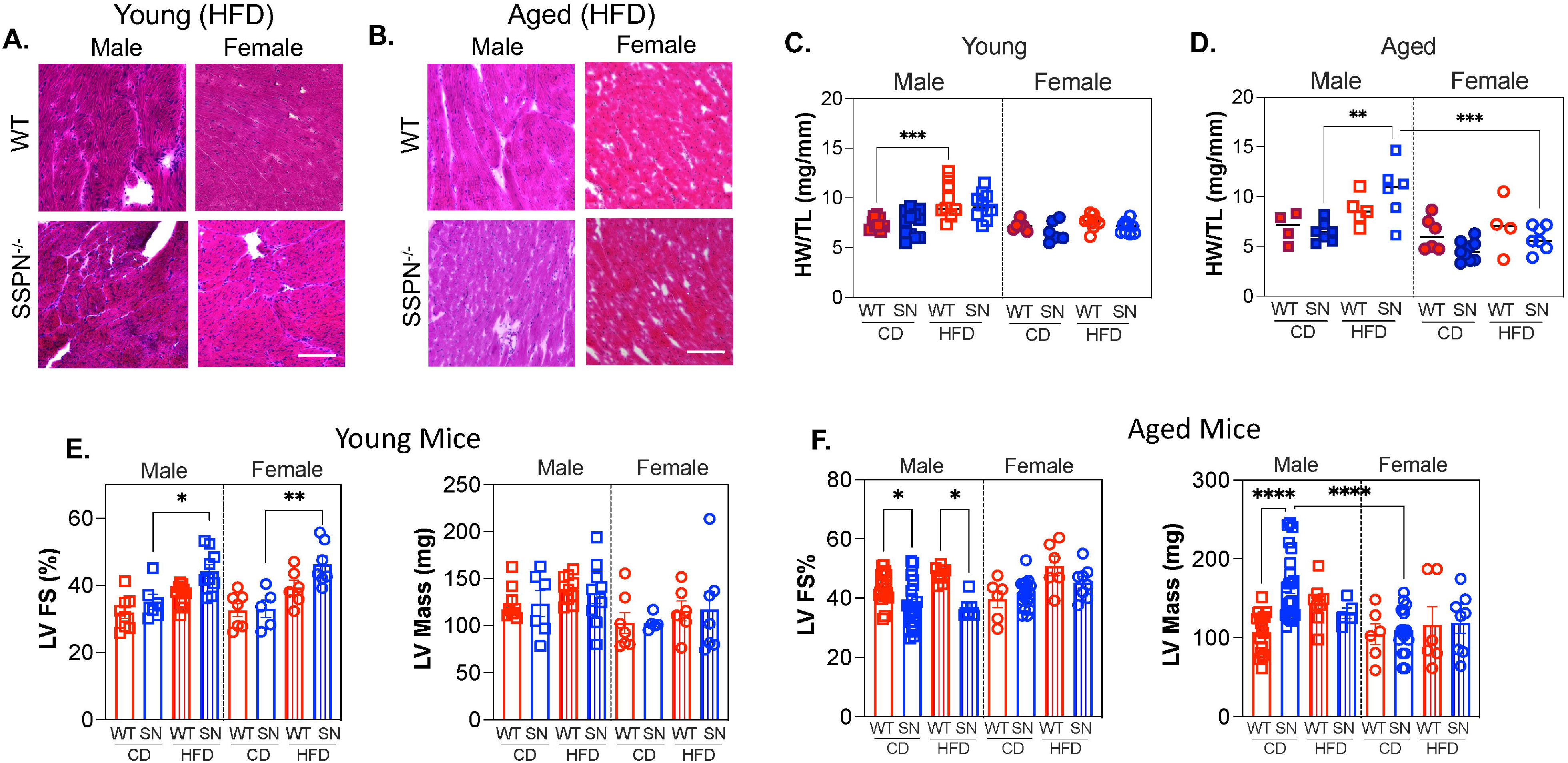
Alterations in heart function in SSPN^-/-^ hearts in response to obesogenic conditions. In **(A)** H& E-stained transverse heart sections obtained from young male and female mice and **(B)** aged male and female mice. Measurements to assess cardiac size heart weight/tibia length (HW/TL) are shown for CD and HFD WT and SSPN^-/-^ (SN) groups in **(C)** young mice and **(D)** aged mice. Echocardiography measurements left ventricular fractional shortening (LV FS%) and LVmass are shown for HFD and CD mice **(E)** young WT and SSPN^-/-^ (SN) mouse groups including **(F)** aged WT and SSPN^-/-^ mouse groups. Numbers of mice used in this study are included in Tables V and VI. Individual values are shown, and statistics were calculated using one-way ANOVA followed by the Tukey’s multiple comparisons post hoc test and *p* ≥ 0.05 considered significant and errors reported as ± S.E.M. and indicated with asterisks, * (<0.05), ** (<0.01), *** (<0.001), **** (<0.0001).

To assess the extent of cardiac remodeling in the face of metabolic stress, HW/TL measurements are shown in Figure 5C for young mice and Figure 5D for aged mice. This provides a crude assessment of cardiac remodeling due to heightened workload due to increased body mass during the diet course. In young mouse groups only WT HFD male mice had a significantly increased HW/TL ratio. Whereas in the aged HFD male mouse groups the trend for increased HW/TL remained for WT mice but significantly increased but variable in SSPN^-/-^ mice. Overall female HFD mice did not exhibit any changes in HW/TL although HW/TL in SSPN^-/-^ females had significantly lower HW/TL than SSPN^-/-^ males. LVmass changes were evident although overall the values in this study were also higher than expected for the CD mice (Figure 5E and 5F). The LVmass was significantly increased in aged SSPN^-/-^ male CD mice although the HW/TL was not increased. After HFD the aged SSPN^-/-^ male mouse group appeared to have ventricular wall remodeling or thinning as the LVmass was lower while the HW/TL was significantly increased (Figure 5F and Table V). Overall, female mouse groups didn’t have any significant changes in LVmass due to obesogenic diet conditions.

**Table V.** Summary of Male Mouse Echocardiography Values in Control and High Fat Diet Studies.

| Young |  |  |  |  | Aged |  |  |  |
| --- | --- | --- | --- | --- | --- | --- | --- | --- |
| ECHO Parameter | WT (CD)<br>(n=8) | SSPN<sup>-/-</sup> (CD)<br>(n=6) | WT (HFD)<br>(n=10) | SSPN<sup>-/-</sup> (HFD)<br>(n=9) | WT (CD)<br>(n=20) | SSPN<sup>-/-</sup> (CD)<br>(n=24) | WT (HFD)<br>(n=9) | SSPN<sup>-/-</sup> (HFD)<br>(n=6) |
| HR (bpm) | 449.9 ± 15.8 | 480.2 ± 18.6 | 460.58 ± 10.4 | 455.15 ± 20.2 | 460.6 ± 10.8 | 449.4 ± 9.9 | 515.3 ± 4.1 | 432.4 ± 12.9 |
| LVID;s (mm) | 2.53 ± 0.13 | 2.40 ± 0.16 | 2.45 ± 0.08 | 1.95 ± 0.12 | 1.85 ± 0.04<sup>a</sup> | 2.12 ± 0.07 | 1.95 ± 0.05 | 2.20 ± 0.08 |
| LVID;d (mm) | 3.69 ± 0.11 | 3.69 ± 0.15 | 3.89 ± 0.10 | 3.47 ± 0.12 | 3.30 ± 0.07 | 3.44 ± 0.09 | 3.73 ± 0.13 | 3.43 ± 0.09 |
| LV;s (ul) | 19.00 ± 3.53 | 20.50 ± 3.32 | 23.05 ± 1.93 | 13.27 ± 2.54 | 10.64 ± 0.62 | 15.47 ± 1.24 | 12.06 ± 0.81 | 16.00 ± 1.70 |
| LV;d (ul) | 54.61 ± 5.02 | 61.41 ± 6.10 | 68.75 ± 5.80 | 56.13 ± 6.53 | 44.55 ± 2.10 | 49.80 ± 2.91 | 59.97 ± 4.57 | 48.94 ± 2.97 |
| LV EF (%) | 60.44 ± 2.58 | 64.98 ± 2.90 | 67.58 ± 1.33 | 75.80 ± 2.28<sup>c</sup> | 75.80 ± 1.25<sup>a</sup> | 68.58 ± 1.86 | 80.29 ± 0.7 | 67.79 ± 1.99<sup>e</sup> |
| LV FS (%) | 31.92 ± 1.81 | 35.14 ± 2.24 | 37.21 ± 1.04 | 44.17 ± 2.56<sup>c</sup> | 43.76 ± 1.14<sup>a</sup> | 38.01 ± 1.54 | 48.14 ± 0.82 | 36.96 ± 1.48 |
| SV (μl) | 34.82 ± 1.63 | 37.57 ± 2.88 | 44.37 ± 2.30 | 37.85 ± 2.68 | 33.21 ± 1.79 | 34.28 ± 2.43 | 47.92 ± 3.93<sup>b</sup> | 32.94 ± 1.34<sup>e</sup> |
| CO (μl) | 14.09 ± 2.27 | 17.82 ± 3.57 | 20.34 ± 1.08 | 17.15 ± 1.43 | 15.14 ± 1.10 | 15.00 ± 1.23 | 24.67 ± 1.95 | 14.77 ± 1.02<sup>e</sup> |
| MV E/A | 1.58 ± 0.29 | 1.47 ± 0.08 | 1.56 ± 0.15 | 1.65 ± 0.19 | 1.23 ± 0.05 | 1.43 ± 0.09 | 1.34 ± 0.32 | 1.33 ± 0.05 |
| IVCT | 15.00 ± 0.01 | 17.16 ± 1.17 | 4.17 ± 0.01 | 15.84 ± 0.01 | 13.69 ± 0.96 | 17.00 ± 1.95 | 6.92 ± 2.75 | 16.42 ± 1.1 |
| LVmass (mg) | 121.8 ± 7.20 | 119.10 ± 16.28 | 111.25 ± 2.92 | 107.96 ± 9.52 | 107.6 ± 5.74 | 165.6 ± 9.23<sup>d</sup> | 144.3 ± 8.27 | 131.1 ± 6.72 |
| Age (mo) | 8.42 ± 0.40 | 6.99 ± 0.24 | 7.22 ± 0.09 | 7.64 ± 0.31 | 18.85 ± 0.68 | 17.20 ± 0.15 | 16.43 ± 0.00 | 17.39 ± 0.10 |
| HW/TL (mg/mm) | 7.45 ± 0.20 | 7.42 ± 0.37 | 9.65 ± 0.56 | 9.21 ± 0.42 | 6.89 ± 0.77 | 6.45 ± 0.39 | 8.68 ± 0.69 | 10.58 ± 1.17<sup>g</sup> |

Definitions: CD (control diet), CO (cardiac output), ECHO (echocardiography), EF % (ejection fraction %), FS % (fractional shortening), HFD (high fat diet), HR, bpm (heart rate, beats per minute), HW/TL (heart weight/tibia length), ID;d (inner dimension; diastole), ID;s (inner dimension; systole), IVCT (intraventricular contraction time), left ventricular (LV), SV (stroke volume), MV E/A (mitral valve E/A), SSPN<sup>-/-</sup> (sarcospan-deficient), WT (wild-type).
One-way ANOVA was used to compare differences between groups. Superscript designations are: <sup>a</sup>, $p < 0.05$ - WT (young vs aged) CD mice; <sup>b</sup>, $p < 0.05$ - WT (young vs aged) HFD mice; <sup>c</sup>, SSPN<sup>-/-</sup> (young) CD vs HFD mice (\*, LV EF%, \*, LV FS%); <sup>d</sup>, $p < 0.05$ - WT vs SSPN<sup>-/-</sup> (aged) CD mice; <sup>e</sup>, $p < 0.05$ - WT vs SSPN<sup>-/-</sup> (aged) HFD mice; <sup>f</sup>, $p < 0.05$ - WT CD vs HFD (aged) mice; <sup>g</sup>, $p < 0.05$ - SSPN<sup>-/-</sup> (aged) CD vs HFD mice.
Significant Groups: WT CD male (young vs aged)(\*\*\*\*, LVID;s, \*\*\*\*, LV EF%, \*\*\*\*, LV FS%, \*\*, CO), HFD WT vs SSPN<sup>-/-</sup> (aged)(\*, CO), CD WT vs KO (aged)(\*\*\*\*, LVmass), WT CD vs HFD (aged)(\*\*, SV), HFD WT (young vs aged)(\*, LV EF%, \*, LV FS%)

Young mice did not show any observable decrements to cardiac function after HFD. The only changes observed were in young male and female SSPN^-/-^ HFD mice, which had significantly increased contractility as evidenced in LV EF% and LV FS% measurements (Figure 5E) and Tables V and VI. With aging, male SSPN^-/-^ CD and HFD mice had a modest but significant reduction in contractility with reduced LV FS% (Figure 5F) and LV EF% (Table V). Aged SSPN^-/-^ HFD mice began showing a reduction in LV contractility measurements compared to young SSPN^-/-^ HFD mice, however their function was still maintained. It may be that the compensatory response to HFD in these mice was starting to have an effect. Values for the contractility measurements were high overall and may reflect low sedation in the mice. Heart rate can affect specific parameters and was largely maintained between 450-500 bpm. The exception however were the aged WT CD male mice with measurements recorded at a higher heart rate, altering these measurements compared to others.

**Table VI.** Summary of Female Mouse Echocardiography Values in Control and High Fat Diet Studies.

| Young |  |  |  |  | Aged |  |  |  |
| --- | --- | --- | --- | --- | --- | --- | --- | --- |
| ECHO Parameter | WT (CD)<br>(n=7) | SSPN<sup>-/-</sup> (CD)<br>(n=5) | WT (HFD)<br>(n=6) | SSPN<sup>-/-</sup> (HFD)<br>(n=7) | WT (CD)<br>(n=6) | SSPN<sup>-/-</sup> (CD)<br>(n=18) | WT (HFD)<br>(n=6) | SSPN<sup>-/-</sup> (HFD)<br>(n=8) |
| HR (bpm) | 461.7 ± 21.1 | 429.9 ± 10.4 | 460.92 ± 11.0 | 445.43 ± 19.7 | 531.8 ± 34.4 | 477.0 ± 12.0 | 509.5 ± 22.0 | 482.2 ± 10.2 |
| LVID;s (mm) | 2.23 ± 0.16 | 2.40 ± 0.22 | 1.80 ± 0.08 | 1.85 ± 0.04 | 1.83 ± 0.20 | 1.95 ± 0.06 | 1.44 ± 0.15 | 1.83 ± 0.09 |
| LVID;d (mm) | 3.44 ± 0.17 | 3.56 ± 0.22 | 3.47 ± 0.18 | 3.28 ± 0.07 | 3.20 ± 0.17 | 3.33 ± 0.06 | 2.88 ± 0.10<sup>a</sup> | 3.34 ± 0.06 |
| LV;s (ul) | 11.62 ± 1.15 | 14.21 ± 1.97 | 16.27 ± 2.16 | 8.96 ± 1.98 | 13.36 ± 3.14 | 12.92 ± 1.15 | 6.89 ± 2.39 | 15.14 ± 1.99 |
| LV;d (ul) | 44.53 ± 3.51 | 48.82 ± 1.65 | 55.06 ± 5.12 | 40.34 ± 2.65 | 41.83 ± 4.86 | 45.68 ± 1.85 | 32.59 ± 3.17<sup>a</sup> | 45.63 ± 1.88 |
| LV EF (%) | 61.74 ± 2.89 | 62.09 ± 3.76 | 70.61 ± 2.56 | 78.18 ± 2.30<sup>c</sup> | 74.98 ± 4.09 | 72.99 ± 1.35 | 82.40 ± 3.30 | 76.6 ± 2.57 |
| LV FS (%) | 32.6 ± 1.98 | 32.99 ± 2.60 | 39.39 ± 2.16 | 46.31 ± 2.37<sup>c</sup> | 39.64 ± 2.80 | 41.47 ± 1.12 | 50.88 ± 3.26 | 45.3 ± 1.96 |
| SV (μl) | 30.41 ± 3.10 | 32.97 ± 3.84 | 36.06 ± 4.71 | 36.59 ± 3.32 | 32.29 ± 2.93 | 32.79 ± 1.01 | 25.71 ± 0.95 | 35.1 ± 0.98 |
| CO (μl) | 12.64 ± 2.17 | 14.30 ± 1.83 | 16.61 ± 2.30 | 14.58 ± 2.07 | 16.18 ± 2.00 | 14.77 ± 1.02 | 12.94 ± 0.71 | 17.0 ± 0.71 |
| MV E/A | 1.97 ± 0.26 | 1.62 ± 0.11 | 2.28 ± 0.19 | 2.27 ± 0.13 | 1.46 ± 0.10 | 1.31 ± 0.19 | 1.34 ± 0.14 | 1.33 ± 0.09<sup>b</sup> |
| IVCT | 13.10 ± 1.20 | 11.85 ± 1.11 | 11.19 ± 0.01 | 11.97 ± 0.01 | 16.18 ± 2.50 | 10.41 ± 0.56 | 13.37 ± 2.18 | 16.42 ± 1.12 |
| LVmass (mg) | 103.0 ± 13.16 | 103.5 ± 4.70 | 108.4 ± 9.51 | 107.6 ± 5.74 | 104.6 ± 13.13 | 109.4 ± 6.60 | 116.1 ± 22.92 | 119.1 ± 13.45 |
| Age (mo) | 8.41 ± 0.40 | 8.35 ± 0.50 | 7.74 ± 0.25 | 8.12 ± 0.51 | 20.20 ± 0.43 | 17.20 ± 0.15 | 18.69 ± 0.40 | 17.17 ± 0.19 |
| HW/TL (mg/mm) | 7.19 ± 0.21 | 6.62 ± 0.42 | 7.77 ± 0.17 | 7.08 ± 0.19 | 6.25 ± 0.68 | 6.34 ± 0.84 | 7.06 ± 1.39 | 6.19 ± 0.41 |

Definitions: CD (control diet), CO (cardiac output), ECHO (echocardiography), EF % (ejection fraction %), FS % (fractional shortening), HFD (high fat diet), HR, bpm (heart rate, beats per minute), HW/TL (heart weight/tibia length), ID;d (inner dimension; diastole), ID;s (inner dimension; systole), IVCT (intraventricular contraction time), left ventricular (LV), SV (stroke volume), MV E/A (mitral valve E/A), SSPN<sup>-/-</sup> (sarcospan-deficient), WT (wild-type).
One-way ANOVA was used to compare differences between groups. Superscript designations are: <sup>a</sup>, *p*<0.05 - WT (young vs aged) HFD mice <sup>b</sup>, *p*<0.05 - SSPN<sup>-/-</sup> (young vs aged) HFD mice (\*, E/A); <sup>c</sup>, SSPN<sup>-/-</sup> (young) CD vs HFD mice (\*, LV EF%, \*, LV FS%)
Significant Groups: Time 0: WT (young vs aged) HFD mice (\* LVID;s, \* LV;d); SSPN<sup>-/-</sup> (young vs aged) HFD mice (\*, E/A); SSPN<sup>-/-</sup> (young) CD vs HFD mice (\*, LV EF%, \*, LV FS%)

Diastolic function measurements were limited in this study. Mild diastolic dysfunction occurs in response to obesity and becomes more severe with increased metabolic dysfunction. The mice in this study ranged from values above 2.0 in young HFD female mice with hyperdynamic cardiac function, indicative of restrictive ventricular filling to more normal values above 1.0. The mice in this study had higher E/A ratio values with very short deceleration time likely due a reduction in ventricular compliance and higher than normal filling pressures (Tables V and VI).

Overall, in young mice the most striking finding was the lower overall fat mass in SSPN^-/-^ females on either CD or HFD. The aging aspect of the study unveiled some genotype and sex-dependent changes with the most profound influences on LVM in aged male SSPN^-/-^ CD mice and glucose tolerance in aged female SSPN^-/-^ CD mice. These findings are supportive of a role for SSPN in influencing glucose handling, affecting fat deposition and perhaps influencing one or multiple factors that drive LVM enlargement.

## DISCUSSION

Dystrophin deficiency has been shown to affect metabolic parameters although the mechanisms are not completely understood. Duchenne/Becker muscular dystrophy (DMD/BMD) patients have been shown to have greater risk of developing insulin resistance, especially deletion carriers at exons 45 and 50 of the dystrophin *DYS1* gene ^39^. To address this muscle fibers from DMD patients were examined and found to have alterations in GLUT4 expression ^39^. The genetic Duchenne muscular dystrophy (DMD) *mdx* mouse model has also been examined to assess how dystrophin deficiency affects metabolic parameters. Strakova et al. reported that *mdx* mice had significantly lower fat mass relative to wild type C57BL/10 mice ^40^. Under thermoneutral conditions *mdx* mice did not exhibit increased fat mass or body weight compared to C57BL/10 mice. However, dystrophin-deficient mice had significant metabolic alterations and altered glucose uptake; suggesting therefore, dystrophin appears to contribute to metabolic regulation ^40^. The sarcoglycan (SG) complex has also been linked to muscular dystrophies accompanied by metabolic defects ^41^.

The findings in *mdx* mice is supported by one of the pathological hallmarks of DGC-related muscular dystrophies - fatty tissue infiltration of skeletal muscle including dystrophinopathies ^42^ and sarcoglycanopathies (LGMD2C-F) ^43^. In muscle diseases that cause atrophic changes or bouts of skeletal muscle degeneration/regeneration the source of fatty infiltrate is activation of adipogenesis-competent cells ^44^. SSPN is abundantly expressed in skeletal and heart muscle ^15, 45^, however, it has been found expressed in other tissue types including ovaries ^15^, subcutaneous adipose tissue (SAT), and omental visceral adipose tissue (OVAT) ^46^. In addition, there is evidence that SSPN impacts metabolic pathways resulting in altered fat distribution ^47^.

Sex differences were identified for a SNP rs10842707 near the *ITPR2-SSPN* gene associated with WHRadjBMI using PAINTOR, a fine mapping tool ^33^. Whereas Keller and colleagues showed strong sexual dimorphism in women for the SNP variant rs718314 that affected DNA methylation of the SSPN promoter, which was associated with WHR (strong sexual dimorphism in women)^18^. Like the human-based studies, our study using SSPN^-/-^ mice also suggested sex differences exist in response to obesogenic diet. Loss of the SSPN protein affected young and aged mice of both sexes and several key similarities were found with the multiple association studies related to anthropometric traits. Limitations exist when utilizing female mice to recapitulate aging effects in humans due to differences in estrogen effects where at equivalent ages women are in menopause as estrogen levels decline after the age of 50 while mice undergo estropause manifesting with enhanced estrogen levels ^48^. After > 13 months of age C57BL6 female mice have decreased cycling however estrogen levels remain similar to young C57BL6 mice ^49, 50^. Therefore, additional measures such as ovariectomy are needed to reduce estrogen levels so aged female mice more closely approximate human female menopause.

SSPN deficiency resulted in lower body weights in mice after HFD administration. Several factors could be contributing to this, including the low-fat content of the standard chow fed (4%) in our study, lower than commonly utilized standard chow diets. Male and female SSPN^-/-^ mice on a mixed (C57BL/6 – 129 SV/J) background fed ad libitum the higher energy concentration NIH-31 modified rodent diet with (5% fat) were found to be heavier than their WT counterparts ^51^. Diet and consumption differences may contribute to lower weight gain seen in SSPN^-/-^ CD mice in our study. The changes in food consumption in aged SSPN^-/-^ mice may be influenced by heightened changes in glutamatergic nerve cell activity that occurs in the lateral hypothalamus of mice fed HFD over a longer time period later in life ^52^ ^53^.

To examine the influence of SSPN on glucose handling mice were administered i.p. glucose to study how sex, metabolic stress, and age affected glucose tolerance in SSPN^-/-^ mice. All the young mouse groups in this study exhibited similar glucose clearance abilities even after 4 months of HFD. Other studies have shown that C57BL/6J mice remain more insulin sensitive than other strains of mice despite higher weight gain in response to diet induced obesity ^54^. An aging study showed that male C57BL/6J mice experience a significant deterioration in glucose tolerance from 6 to 18 months of age not observed in female mice^55^. In contrast our study used mice at 6 and 18 months of age, and we did not observe significant changes in the AUC of aged male WT mice. This difference may be due to the glucose dose (1 g/kg) in our study compared to 2g/kg in the previously referenced study. By the end of the CD regimen aged SSPN^-/-^ male and female mice had lower % fat (Figure 3A and 3B) and similar lean body mass as WT but poorer glucose clearance (AUC). This suggests that the SSPN^-/-^ mice are more metabolically unhealthy than the WT mice with a poor tissue response to insulin. In human association studies the *SSPN* locus was identified with strongest associations with increased waist-hip-ratio (WHR) ^56^ and a SNP associated with increased waist-height relationship adjusted body mass index (WHRadjBMI), fasting insulin adjusted for body mass index (FIadjBMI) and type 2 diabetes ^10^. Loss of SSPN in mice protected them from fat deposition and weight gain under controlled diet and activity. On a low-fat standard chow diet, the aged SSPN mice developed glucose intolerance compared to WT mice, therefore other factors must be contributing to insulin resistance independent of weight gain. From this finding it is anticipated that SSPN^-/-^ mice may be more susceptible to type 2 diabetes while humans with specific SNPs in the SSPN locus may have enhanced risk through increased central adiposity.

The glucose intolerance seen in the lean CD SSPN^-/-^ mice provides an opportunity to examine pathophysiological mechanisms related to SSPN deficiency compared to that of “lean” diabetes. SSPN deficiency combined with ageing led to glucose intolerance and fasting blood glucose levels in the diabetic range for female CD and male and female HFD, which were leaner than their WT counterparts. In lean diabetes rapid beta cell failure causes impaired pancreatic insulin secretion, which is more likely to drive hyperglycemia than insulin resistance ^57^. The lower weights of young 2-month-old SSPN^-/-^ compared to WT female correlate to lower birth weights supporting a link demonstrated between low birth weight, lean individuals and development of type 2 diabetes in lean individuals ^58, 59^. Accompanying physiological defects were observed in these individuals including reduced insulin secretion, muscle glucose uptake, and insulin stimulated glycolysis ^58, 59^. Initial studies were performed in young mice and no overt phenotypic change was found in their ability to clear glucose during GTT.

Aging however influenced the phenotype and SSPN^-/-^ mice had noticeable alterations in glucose handling. The aged CD SSPN^-/-^ females had significantly higher AUC compared while aging interfered with glucose clearance response of all the HFD mice. The reduction in glucose tolerance of aged female SSPN^-/-^ mice may be the result of several changes including increased VAT mass compared to younger female SSPN^-/-^, however the % Fat was largely unchanged. This could indicate changes in distribution from subcutaneous to visceral fat pad storage. Multiple studies have shown that intraabdominal or visceral adipose tissue storage is a major contributor to metabolic risk including insulin resistance ^60^ ^61^ while others suggest a protective role for subcutaneous adipose tissue ^62^.

In this study age may play a role by influencing the levels of freely circulating sex hormones, which influences adipose storage. In HFD studies in C57BL/6J mice both male and ovariectomized female mice have been found to store more abdominal adipose, perhaps in the form of hypertrophied adipocytes ^63^. Estrogen affects fat distribution in females by influencing insulin-sensitive glucose uptake. In addition, estrogen strongly inhibits key adipogenic genes at the mRNA level including leptin and hormone-sensitive lipase (HSL). Studies in younger female mice have shown that a significant correlation exists between small adipocyte size and insulin sensitivity ^64^. In our study, however, aged HFD SSPN^-/-^ female mice had significantly larger adipocytes than WT but not large enough to be considered hypertrophic. This may reflect age-related changes that can be attributed to the functional decline of adipocyte progenitors and senescent cell accumulation ^65^. Capacity for adipogenesis declines by late middle age in humans and experimental animal models with a reduction of peroxisome proliferator-activated receptor gamma (PPARy) and CCAAT/enhancer-binding protein alpha (C/EBPa) expression is accompanied by decreased adipose tissue mass and metabolic function ^66^. Young SSPN^-/-^ mice had modest alterations in % Fat and VAT weight after 4 months HFD had little alteration in adipocyte size as seen in WT HFD female mice. Adipocyte hyperplasia is the most favorable means of adipose tissue expansion and provides protection against metabolic disease since it allows lipid storage and preserves normal adipocyte function. Development of large adipocytes as seen in male HFD WT and SSPN^-/-^ VAT can lead to increased production of hormones and bioactive substances including leptin ^67^ and insulin, pro-inflammatory cytokines and reactive oxygen species (ROS) ^68^.

To better understand how the alterations in adipose deposition and distribution in SSPN^-/-^ mice might impact cardiac performance, cardiac function was assessed by echocardiography. Previous studies in mice have demonstrated that HFD-induced obesity and hyperglycemia are not always sufficient to induce cardiac dysfunction ^69^. The adult mouse heart is highly adaptable and adjusts to sustained levels of high intracellular glucose by switching to a fetal-like metabolic pattern for long periods without adverse effects on cardiac function ^70^. Increased fat in the diet leads to increased fatty acid oxidation and increased oxidative stress, however if the metabolic flexibility of the heart is impaired it may increase oxidative stress. SSPN deficiency has been shown to decrease the oxidative capacity of cardiomyocytes, however this has not been studied in vivo ^38^.

Overall, in our studies with young and aged mice we did not find any decrements to cardiac function. Young male and female HFD SSPN^-/-^ mice exhibited signs of hyperdynamic cardiac function with increased LV EF% and LV FS% compared to CD controls. The only significant alteration in young mice in response to HFD was increased HW/TL in the male WT mice. Cardiac hypertrophy in response to HFD has been well documented in C57BL/6J mice ^71^ and in this study it was greatest in young male WT HFD mice, which also exhibited the greatest body mass increase. LVmass was significantly higher in aged SSPN^-/-^ male mice under baseline CD conditions.

LV mass was only slightly altered by HFD in aged WT mice while aged male SSPN^-/-^ hearts had significantly higher LV mass at baseline. Doppler echocardiography was used to assess changes in diastolic function. In young mice, the largest alterations in E/A ratio, the ratio between early and late ventricular filling, was observed in female HFD mice. This suggests an increase in LV passive stiffness that results in most of the blood entering the ventricles in an early filling pattern with abrupt termination of filling. The E/A ratios in the aged mice were reduced compared to young but still in the normal range suggesting mild to no diastolic dysfunction.

Several cardiac structural parameters were altered in aged male SSPN^-/-^ mice as they had significantly higher baseline LVmass values than matched WT controls. Aged SSPN^-/-^ HFD mice however, showed signs of ventricular remodeling with reduced LVmass compared to SSPN^-/-^ CD and WT HFD mice. The HW/TL values were significantly larger in the aged male HFD SSPN^-/-^ mice compared to their respective control. Since cardiac function was largely preserved and there were no indications of chamber remodeling it appears that these mice were still maintaining compensatory function. This supports earlier studies that indicate that the SSPN locus plays a distinct role in determining cardiac mass ^24^. Future studies will be directed towards understanding factors that may contribute to this phenotype in aging SSPN^-/-^ male mice and how obesogenic conditions affect cardiac remodeling and function.

## ACKNOWLEDGEMENTS

The authors thank the following agencies for support during these studies. M.S.P. received funding from Florida Department of Health James and Esther King Foundation #21K21, FSU FYAP, and AHA Scientist Development Grant 16SDG29120002. The authors have no conflicts to disclose. The content here within is solely the author’s responsibility and does not necessarily represent the official views of the Florida State University.

## AUTHOR CONTRIBUTIONS

**ARK** (aged mice echocardiographic measurements and analysis, weight and food consumption checks, body composition measurements, glucose tolerance testing, DNA methylation, figure assembly), **ICV** (young mice echocardiographic measurements and analysis, weight and food consumption checks, glucose tolerance testing), **LS** (aged mice glucose tolerance testing, WAT imaging, adipocyte measurements, table assembly), **RQC** (WAT imaging, tissue histology, adipocyte measurements), **SE** (histology and tissue collection), **RTK** (echocardiography analysis), **BO** (aged mice glucose tolerance testing and body composition measurements), **ARM** (young mice body composition measurements), NM (tissue histology, DNA methylation), **MSP** (study design, data analysis, figure and table assembly, manuscript preparation and writing).

## FIGURE LEGENDS

**Supplemental Figure 1.** In **(A)** area under the curve (AUC) was calculated from daily feed consumption by young HFD mice over the course of the 4-month feeding regimen. **(B)** AUC is shown of daily food consumption by aged HFD mice over the course of the 5-month feeding regimen.

## Notes

### Competing Interest Statement

The authors have declared no competing interest.

