## Supplementary figures and images for "Evaluating the Sex Dependent Influence of Sarcospan on Cardiometabolic Disease Traits in Mice"

### Rahimi Kahmini Suppl Figure 1-v2.pdf

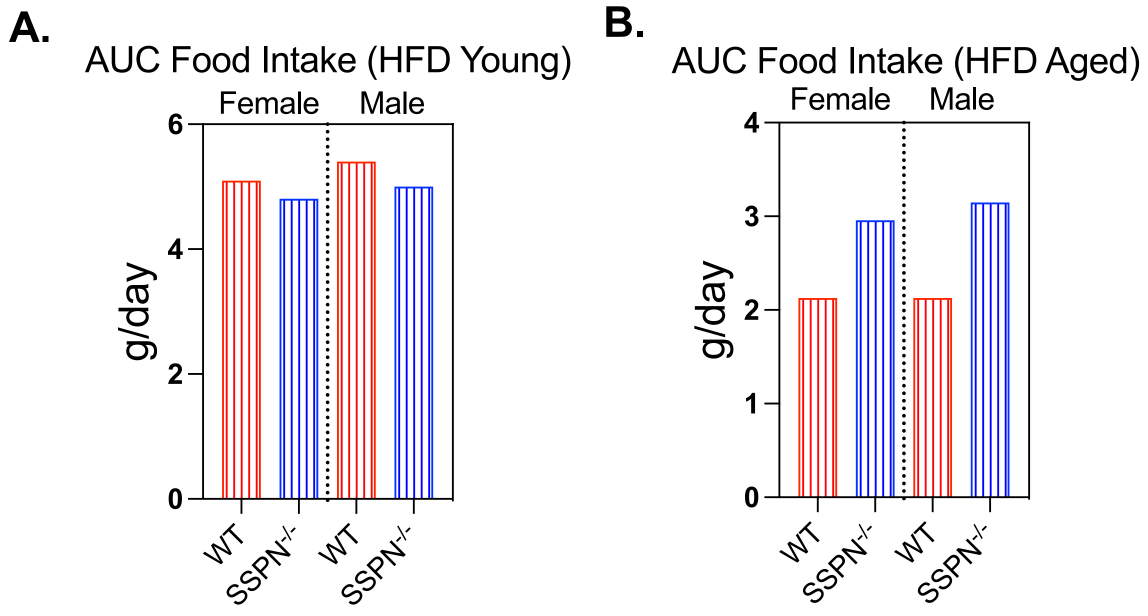


Rahimi Kahmini et al. 2024
Supplemental Figure 1
